# Target Control in Logical Models Using the Domain of Influence of Nodes

**DOI:** 10.1101/243246

**Authors:** Gang Yang, Jorge G. T. Zañudo, Réka Albert

## Abstract

Dynamical models of biomolecular networks are successfully used to understand the mechanisms underlying complex diseases and to design therapeutic strategies. Network control, and its special case of target control, is a promising avenue toward developing disease therapies. In target control it is assumed that a small subset of nodes is most relevant to the system’s state and the goal is to drive the target nodes into their desired states. An example of target control would be driving a cell to commit to apoptosis (programmed cell death). From the experimental perspective, gene knockout, pharmacological inhibition of proteins and providing sustained external signals are among practical intervention techniques. We identify methodologies to use the stabilizing effect of sustained interventions for target control in logical models of biomolecular networks. Specifically, we define the domain of influence of a node (in a certain state) to be the nodes (and their corresponding states) that will be ultimately stabilized by the sustained state of this node regardless of the initial state of the system. We also define the related concept of the logical domain of influence of a node, and develop an algorithm for its identification using an auxiliary network that incorporates the regulatory logic. This way a solution to the target control problem is a set of nodes whose domain of influence can cover the desired target node states. We perform greedy randomized adaptive search in state space to find such solutions. We apply our strategy to several biological networks to demonstrate its effectiveness.

## 1 INTRODUCTION

In cellular systems various molecular species, such as DNA, RNA, proteins and small molecules, interact in diverse ways. The totality of these interactions gives rise to cellular functions. The relationship between molecular interacting systems and cellular functions is studied in the new emerging field of systems biology [1, 2]. A promising systems biology methodology is to represent the molecular interacting system as a network, construct a dynamic model of the information propagation on this network, and identify the cellular functions with long-term behaviors of the dynamic model [2, 3, 4, 5]. Various dynamical models of biological networks have been built to integrate related experimental results and to reveal the underlying mechanisms of complex diseases such as cancers. Among various types of dynamical models, logical models, such as Boolean network models, have the advantage of being scalable and not requiring detailed knowledge of kinetic parameters [5, 6]. An abundance of recent literature has shown that logical models can capture the emergent behaviors of real biological systems, they can generate predictions that are validated by follow-up experiments and they can predict successful intervention strategies [7, 8, 9, 10, 11].

Network control has recently become a popular research topic as it reflects our interest to not only understand an interacting system, but also intervene in it and modify its outcomes [12, 13]. Network control is a broad subject; different underlying models, different control goals and different possible interventions can be considered [12]. Various control strategies have been designed for both continuous dynamical systems [14, 15, 16, 17, 18, 19] and discrete ones [20, 21, 22, 23]. In electric circuits modeled by a system of linear ordinary differential equations, it is possible to obtain full control of the system, that is, to drive the system to any state from any initial condition [24, 14]. For non-linear systems, attractor control, that is to drive the system to one of its natural attractors from any initial condition, has been achieved in several modeling frameworks, such as feedback vertex control for ordinary differential equation models [18] and stable motif control for logic (Boolean) models [20]. However, in biological systems it is not necessary and often not practical to control every component of the system. A more realistic problem is target control, where we assume that the state of the system is mostly determined through a subset of nodes and the control goal is to drive these nodes into desired states. The target control problem has been considered for continuous models by [25]; here we provide a framework to solve the target control problem in Boolean network models.

Despite recent progress in molecular biology, quantitatively manipulating the level of a chemical species is still a challenging problem for experimentalists. Thus any control strategy involving applying time-dependent, variable signals to a target is hard to implement in real systems. However, gene knockout, pharmacological inhibition of proteins and providing sustained external signals have been robustly implemented and demonstrated to be effective intervention strategies [26]. Thus we choose our intervention options to be maintaining a sustained state (either absence or abundant activity) in order to make the solution more practical. To describe the long-term effect of such a sustained state, we define a node property called domain of influence for each node. This is essentially asking which other nodes will adopt a fixed state due to the sustained intervention regardless of initial conditions. Then it follows that the solution to the target control problem will be the set(s) of nodes whose domain of influence can cover the desired target node state combinations.

In the following, we first introduce the Boolean modeling framework and relevant previously-developed concepts such as the expanded network and stable motif. Then we define the domain of influence (DOI) and logical domain of influence (LDOI) of a node or multiple nodes and present several useful properties. We then define our target control problem and describe our target control strategy based on DOI using greedy randomized adaptive search in state space. We finally illustrate the effectiveness of our target control strategy in random ensembles and real biological network models.

## 2 MATERIALS AND METHODS

### 2.1 Boolean network models of biological systems

A dynamical model of a biological system starts with the construction of a network consisting of nodes that represent the system’s elements and edges that specify the pairwise relationships between nodes. In biological networks at the molecular level, nodes are molecular species such as small molecules, RNA, protein, and edges indicate interactions and regulatory relationships. In discrete models, each node i is characterized by a discrete state variable *σ_i_*, and the vector (*σ*_1_,…, *σ_n_*) represents the state of the system [7]. The state of the system can be followed in continuous time or at discrete time intervals. In discrete time models, each node state is updated at discrete time intervals. Mathematically, the activity of each node *σ_i_* is described by a regulatory rule 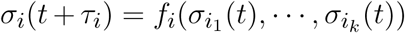, where *i_1_*,…, *i_k_* are the regulating nodes of *i* and *τ_i_* is a discrete time delay [7]. The regulatory functions *f* cannot be constant functions (i.e. cannot yield the same output regardless of the state of the regulators). In models describing signal transduction networks the external signals are represented with source nodes whose regulatory functions depend only on their own state, usually sustaining this state: *σ_i_*(*t* + *τ_i_*)= *σ_i_*(*t*) [7].

Here we focus on discrete time Boolean network models, where node states are binary, 1(ON) or 0(OFF), and the regulatory function is specified by a truth table or using the Boolean operators AND, OR, NOT [27, 28, 7]. This is motivated by the fact that biological species are frequently observed to demonstrate switch-like behaviors and have highly nonlinear regulations; thus the node state 1 means the molecular species is above a threshold concentration or activity and thus it is able to regulate its targets and the node state 0 means it is below a threshold concentration or activity and is thus ineffective [29, 5]. Depending on the updating scheme, the time trajectory of the system is simulated deterministically or stochastically. A simple deterministic updating scheme is synchronous updating, where *τ_i_* =1 for every node [5]. In this scheme, the system will deterministically evolve from a specific initial state into an attractor, which can be a steady state (fixed point) or a limit cycle, which consists of several states that repeat regularly. Steady states can be interpreted as cell types; limit cycles may correspond to a cell cycle or circadian rhythms [7]. In general asynchronous updating, a commonly used stochastic updating scheme, a random node is selected to be updated at each time step [30]. This type of update is motivated by the fact that different biological processes have various timescales, and often the timescales of specific processes are not known [31]. While limit cycles depend on the specific chosen updating regime, fixed points (steady states) do not depend on the updating scheme [32]. Stochastic update may lead to attractors that involve irregular repetition of a set of states, called complex attractors.

### 2.2 Previously established concepts in Boolean networks

The expanded network integrates the original network with the regulatory rules of each node [33]. We illustrate the expanded network with the example in Fig. 1, which consists of five nodes, node 0, 1, 2, 3 and 4 with the regulatory functions *f*_0_ = NOT *σ*_3_, *f*_1_ =(NOT *σ*_0_) OR *σ*_3_, *f*_2_ = NOT *σ*_1_, *f*_3_ = (NOT *σ*_2_) OR (NOT *σ*_4_), *f*_4_ = *σ*_0_ OR *σ*_1_. First, we denote each original node i by *n_i_* in the expanded network, and we introduce a complementary node for each original node in the system to represent the negation (deactivation) of the original node, denoted by ~*n_i_* [33]. As the NOT function is a unary operator, all the NOT functions are replaced by the negated state of the respective node (*i.e.* its complementary node) in each Boolean regulatory function. The edges in the expanded network are revised according to the updated rules so that every edge represents a positive regulatory relationship in the expanded network. For example, *f*_0_ = NOT *σ*_3_ implies the rule for the original node *n*_0_ as 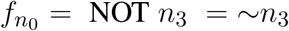, and thus a corresponding edge is drawn from ~*n*_3_ to *n*_0_ in the expanded network. The Boolean regulatory function for the complementary (negated) node is the logical negation of the regulatory function of the original node. In this example, 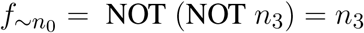 and thus a corresponding edge is drawn from *n*_3_ to ~*n*_0_ in the expanded network.

**Figure 1.**
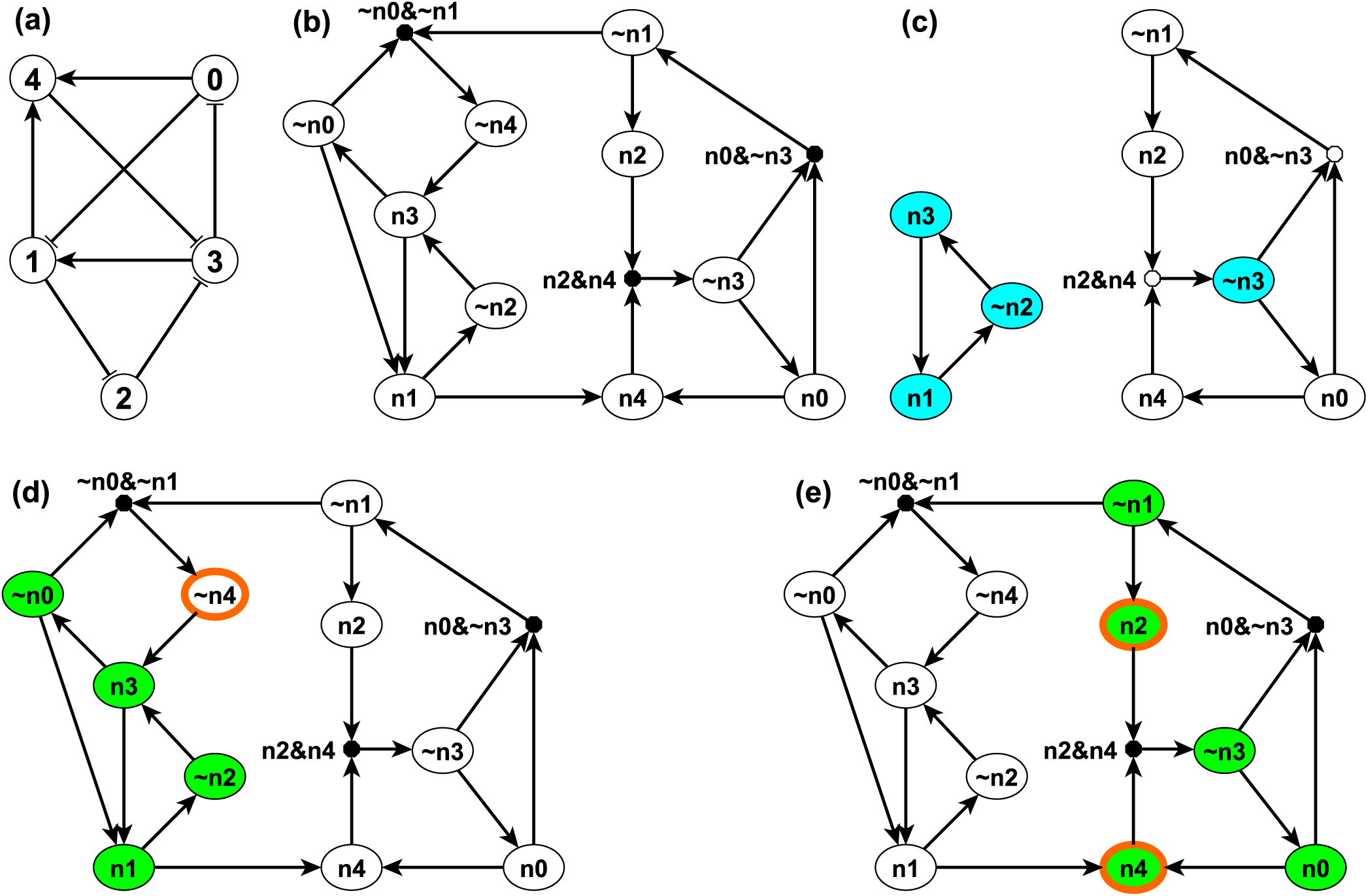
An example network, its corresponding expanded network and its stable motifs are shown in sub-figures (a), (b) and (c) respectively. The LDOI of {~*n*_4_} and {*n*_2_, *n*_4_} illustrated on the expanded network are shown in sub-figures (d) and (e) respectively. In panel (a) each edge with an arrow represents activation and each edge with a flat bar represents inhibition. Each node i in panel (a) has a correspondent *n_i_* and its complementary node ~*n_i_* in panel (b). (Note that *n_i_* is labeled as *ni* in panel (b) to be more visible). A composite node is drawn as a filled black circle and & represents the AND logic operator. In panel (c), each blue node is a single-node core of the corresponding stable motif. In panel (d) and (e), nodes with thick orange boundary are the sustained interventions and the green nodes are their LDOI.

Second, to differentiate OR rules from AND rules when multiple edges point toward the same target node, we introduce a composite node for each set of edges that are linked by an AND function [33]. In order to uniquely determine the edges of the expanded network, the regulatory functions need to be specified in disjunctive normal form, that is, a disjunction of conjunctive clauses (in other words, grouped AND clauses separated by OR clauses). For example, (*A* AND *B*) OR (*A* AND *C*) is in a disjunctive normal form, while *A* AND (*B* OR *C*) is not. The desired disjunctive normal form can be formed by a disjunction of all conditions that give output 1 in the Boolean table and then simplified to the disjunction of prime implicants (Blake canonical form) by the Quine-McCluskey algorithm [34]. Now we add a composite node for each AND clause in the Boolean regulatory function, denoted by a filled black circle in Fig. 1 (b). For example, the composite node ~*n*_0_&~*n*_1_ in the left upper part of Fig. 1 (b) represents the expression (NOT *n*_0_) AND (NOT *n*_1_), which implies the complementary node ~*n*_4_. Notice that one can read all the regulatory functions from the topology of the expanded network. The AND rule is indicated by a composite node with multiple regulators, while all the other edges represent independent activation (parts of an OR function).

As the expanded network encapsulates the regulatory logic that determines the network dynamics, it can serve as a basis for attractor analysis. One approach is through analyzing the stable motifs of the expanded network [35]. A stable motif is defined as the smallest strongly connected component (SCC) satisfying the following two properties: 1) The SCC cannot contain both a node and its complementary node and 2) If the SCC contains a composite node, it must also contain all of its input nodes [35]. The first requirement guarantees that the SCC does not contain any conflict in node states and the second requirement guarantees that all the conditional dependence is satisfied and the SCC is self-sufficient in maintaining each node state inside the stable motif. Thus the stable motif represents a group of nodes that can sustain their states irrespective of outside nodes’ states. The corresponding node states implied by the stable motif can be directly read out: an original node represents the ON (1) state and a complementary node represents the OFF (0) state [35]. For example, in the left part of Fig. 1 (c), node *n*_1_, ~*n*_2_ and *n*_3_ form a stable motif, representing that node 1 and node 3 are ON and node 2 is OFF. There is a strong correspondence between stable motifs and the attractors of the system. Specifically, there is a one-to-one correspondence between a sequence of stable motifs and a fixed point or a partial fixed point (a part of a complex attractor). A partial fixed point is defined as a true subset of all the nodes whose respective state remains unchanged after being updated regardless of the states of the nodes excluded from this subset [35].

### 2.3 The domain of influence of a sustained node state

We define the domain of influence (DOI) of an intervention that maintains a sustained node state as all the node states that will be stabilized in the long term under the influence of this intervention for all initial conditions in any updating regime. Mathematically, 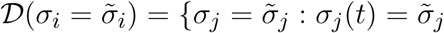 *as t* → ∞ *for any* (*σ*_1_(*t* = 0),…, *σ_k_*(*t* = 0)) *when* 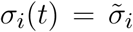 *for any t* > 0}, where 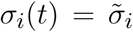 is the intervention, 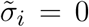 represents knockout and 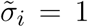 represents sustained activation, 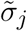 represents a node state fixed by the intervention, and (*σ*_1_(*t* = 0),…, *σ_k_*(*t* = 0)) represents the initial condition of all the nodes of the system. We do not include the intervention node state 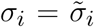 in its own DOI, unless the node is sufficient to maintain the corresponding node state in the long term even in the absence of a sustained intervention. Notice that there is one-to-one correspondence between a node state 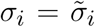 and a non-composite node *n^ex^* in the expanded network: *σ_i_* =1 corresponds to a normal node *n_i_* in the expanded network and *σ_i_* = 0 corresponds to a negation node ~*n_i_*. Thus we use the two notations interchangeably, that is, 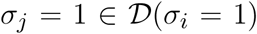 is equivalent to 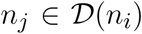 and 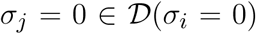 is equivalent to 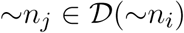.

The DOI of a node is difficult to calculate because it entails determining the common part of all attractors of a dynamical system to identify the nodes whose states stabilize due to the considered intervention. As an alternative to this computationally hard problem, we define a related concept called the logic domain of influence (LDOI) of an intervention that maintains a sustained node state. The logic domain of influence consists of all the node states that, for any initial condition, are stabilized by the first update of the corresponding node in an updating regime that preserves the level order (breadth first search order) of the expanded network. An updating regime preserves the level order if all the nodes in the *n*th level are updated at least once before updating any node in the (*n* + 1)th level (see details in Supplementary Material 2.1). We denote the LDOI of a node state *σ_i_* as 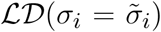. We define the LDOI of an empty set to be an empty set, 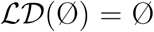. This is consistent with the definition as an updating order preserving the level order starting from a null set can start from any node, and a node will not be stabilized to a fixed state upon its very first update for all initial conditions unless its regulatory function is a constant. Source nodes stabilize in their initial state, which nevertheless will be different for different initial conditions.

### 2.4 Determining the logical domain of influence of a sustained node state

We propose to find the LDOI of a node state by doing a modified breadth first search (BFS) on the expanded network (see the pseudocode in *Supplementary Material* Sec. **1.1**). In order to find the LDOI of 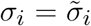, we start the search from the corresponding node (or complementary node) on the expanded network. If we meet another non-composite node, we add this node to the LDOI; if we meet a composite node, we add this composite node only if all of its parent nodes (i.e. regulators) are already part of the LDOI. This is due to the fact that any edge from a node to a non-composite node represents a sufficient relationship and any edge from a node to a composite node represents a necessary relationship. We keep searching on the expanded network until no new nodes can be added to the LDOI. For example, in Fig. 1 (b), one can readily see that 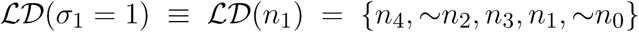 following the described search procedure. The first difference from a normal BFS to find a connected component starting from a node is that we put an extra rule for including a composite node. Another subtle difference is that we do not include the starting point unless we visit this starting point again in our search process.

During the search process, there is a possibility that we meet the negation of the starting point. This reflects the possibility that a node state can indirectly lead to the opposite state through a negative feedback loop. This outcome represents a conflict with the original intervention. We do not add this node to the LDOI because we assume that the intervention can sustain the original node state, thus the opposite state is not reachable. This truncation of the LDOI to avoid including the negation of the starting node state ensures that the LDOI will not contain a node which is the negation of an already visited node. Mathematically, if a non-composite node 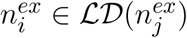, then 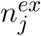 is sufficient to activate 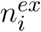, i.e., the long-term logical rule for 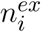 can be expressed in the form 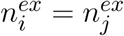; this implies 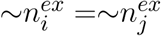 AND…, 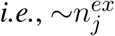 is necessary to activate 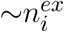. Thus any conflict between 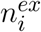 and 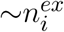 will occur after the conflict between 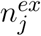 and 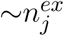 during the search process. This truncation of the LDOI is the third difference compared with a normal BFS.

For example, in the network of Fig. 1 (d), the LDOI of the complementary node ~*n*_4_ includes nodes *n*_3_, ~*n*_0_, *n*_1_, ~*n*_2_ following the search procedure. From *n*_1_ one can also reach node *n*_4_, which is the negation of the considered intervention. Thus we stopped this branch of searching based on our truncation rule. Since there are no more nodes that can be added, we conclude that 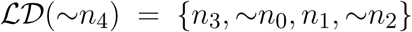.

Our LDOI search procedure is equivalent to doing a simulation on the expanded network. If we update the system corresponding to the BFS order of the expanded network starting from the intervention node (i.e., we update node *i* if we visited 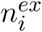 on the expanded network), all the updated nodes are guaranteed to stabilize in the corresponding visited state on the expanded network, *i.e.* as in the logic domain of influence (LDOI) of that node. In the example of Fig. 1, as discussed above, 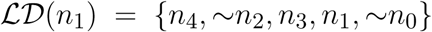. If we update the nodes in the order 4, 2, 3, 1, 0, each node will stabilize in the state as in 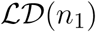. We note that this does not put a restriction on the updating regime: if we update the system in an arbitrary order, each node in the LDOI of the given sustained intervention will stabilize in the first update after all of its regulators included in the LDOI have been updated once. For example, if we fixed the node 1 to be ON and we perform rounds of update of the nodes in the order 0, 1, 2, 3, 4, nodes 2, 3 and 4 will be stabilized in the first round of updating, while nodes 0 and 1 will be stabilized in the second round.

The difference between the LDOI and DOI is that LDOI requires the nodes to be stabilized when being updated for the first time, while DOI just requires the nodes to be stabilized in finite time. Thus one can see that the LDOI of a node will be a subset of the DOI of a node. In many cases the two concepts give the same result. Two exceptions are illustrated in Fig. 2. In both cases the DOI of an intervention contains more nodes than the LDOI of this intervention. This is because certain nodes may stabilize not because of the influence of the intervention but because of the collective effect of two inconsistent feedback loops or because of a stable motif stabilized by an oscillation. In the network of Fig. 2 (a), the three regulators of node B are independent and the network includes both a positive and a negative feedback loop. To analyze the LDOI of *A* =1, taking the feedback effect of C and D on B into consideration, the regulatory function of B is simplified into *σ_B_*(*t* + *τ_B_*)= *σ_B_*(*t* − *τ_C_*) OR NOT (*σ_B_*(*t* − *τ_D_*)), which yields a constant state *σ_B_* =1. Thus 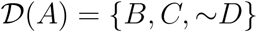, as the stabilization of B leads to the stabilization of C and D as well. However, 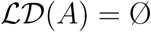 as the activation of the composite node requires nodes *A*, ~*C*, ~*D* on the expanded network shown in Fig. 2 (b) and thus we cannot add the composite node to the LDOI of node A. In the example shown in Fig. 2 (c), the two regulators are independent for node B, 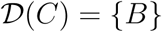 as the negative feedback loop of node A will make A oscillate, but B will stabilize into the ON state after the first time that A visits the ON state and activates B, while 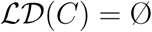 for the same reason as in the last example.

**Figure 2.**
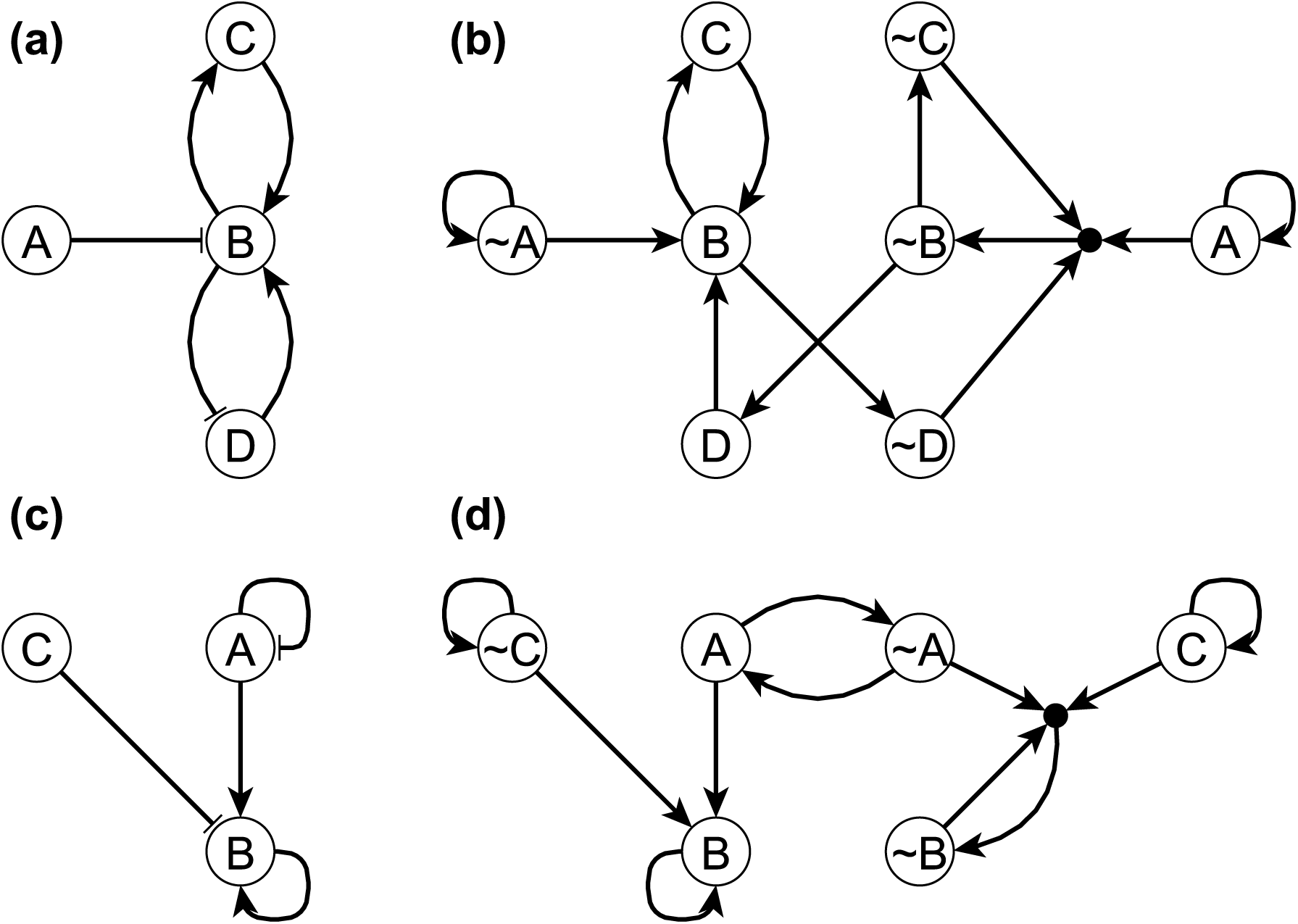
Two example networks (panel (a) and (c)) and their respective expanded networks (panel (b) and (d)) that illustrate the difference between DOI and LDOI. In both networks, an edge with an arrowhead represents activation while an edge with flat bar represents inhibition. Implicit positive self-loops for source nodes are not shown in panel (a) and (c). In panel (a) the regulatory functions are *f_A_* = *A*, *f_B_* =(NOT *A*) OR *C* OR *D*, *f_C_* = *B*, *f_D_* = NOT *B*. When A=0 the system has a single attractor, the fixed point is *σ_B_* = *σ_C_* = 1, *σ_D_* = 0. In panel (c) the regulatory functions are *f_A_* = NOT *A*, *f_B_* = *A* OR *B* OR NOT *C*, *f_C_* = *C.* When *σ_C_* =1 the system has a complex attractor in which A oscillates and *σ_B_* = 1.

### 2.5 Properties of the logical domain of influence of a sustained node state

In order to further illustrate the concept of LDOI, we discuss a few of its properties and its relationship with established concepts in Boolean dynamics.

A natural question to ask is about the possible inclusion relationship between the logic domains of influence of two node states 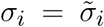 and 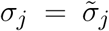 in the case when 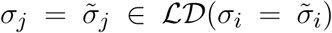 or 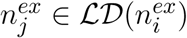 in the expanded network notation, where 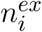 and 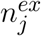 represent any non-composite node in the expanded network. In a directed graph, if node *n_j_* is a reachable from node *n_i_*, all descendants of *n_j_* will also be reachable from *n_i_*; indeed one can easily prove this by contradiction. However, due to the special properties of the expanded network and the truncation of the LDOI, this inclusion relationship 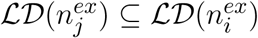 is not generally true for the expanded network. It is possible that 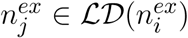, however, 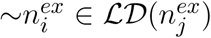. In this case, by definition of the logic domain of influence, we won’t allow the negation of a node state to be part of the logic domain of influence of a node state. For example, 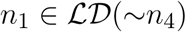, however, 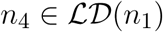. Thus 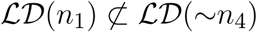.

If we add an additional restriction on the two nodes, this inclusion relationship will hold the same way as for descendants in a directed graph. To be specific, the *first key property of the LDOI* is, if the node state 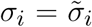 and 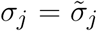, corresponding to the two non-composite node 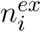 and 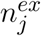 on the expanded network, are both included in the same (partial) fixed point and 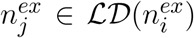, the logic domain of influence of 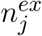 will be a subset of the logic domain of influence of 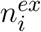, i.e. 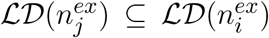. (Recall that a partial fixed point is a subset of nodes whose respective state remains unchanged after being updated regardless of the states of the nodes excluded from this subset.) The reason why the inclusion relationship holds is that node states in a (partial) fixed point stabilize in the long term, thus they will not lead to a situation with opposing behavior 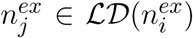 and 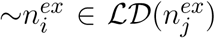. This restriction can be weakened to only require that node state 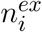 is in a (partial) fixed point. The reason is that if 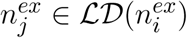 and 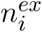 is in a (partial) fixed point, then 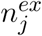 must also be in the same (partial) fixed point, or be a node whose state stabilizes due to the nodes in the partial fixed point. Also, as one or more stable motifs are part of a (partial) fixed point, the conclusion will be true if one replaces “(partial) fixed point” by “stable motif” in the above statement. For example, as nodes *n*_1_, ~*n*_2_ and *n*_3_ form a stable motif and its corresponding (partial) fixed point is (*σ*_1_,*σ*_2_,*σ*_3_) = (1, 0, 1), which also lead to the stabilization of the remaining two nodes as *σ*_0_ = 0 and *σ*_4_ = 1, thus 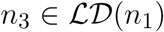 implies that 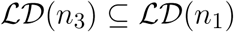. In fact, 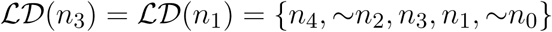. Also 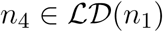 implies that 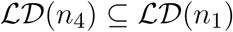. Note that only *n*_1_ is part of the stable motif or partial fixed point in the latter example, *n*_4_ is not.

As stable motifs represent generalized positive feedback loops of the Boolean network [35], we explore the relationship between stable motifs and the logic domain of influence of a node state. The *second key property of LDOI* is, if the logic domain of influence of a node state contains this node state itself, the logic domain of influence contains a stable motif. As the LDOI of a node state only contains the node state itself if we meet this node during the search process on the expanded network, this indicates the existence of a positive feedback loop, which is the intuition why this proposition holds. (A sketch of proof from the dynamical standpoint is included in *Supplementary Material* Sec. **2.2**.) For example, 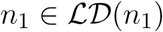 implies that there exists a stable motif contained in 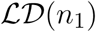, indeed, 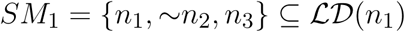.

### 2.6 The domain of influence of a node state set

Now we generalize the concept of DOI of a single node state to DOI of a node state set (i.e., a set of nodes, each in a sustained state). We define the DOI of a node state set as all the node states that can be stabilized in the long term by the given set of node states under all initial conditions in any updating regime. Mathematically, 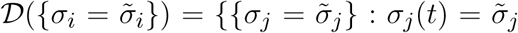 *as t* →∞ *for any* (*σ*_1_(*t* = 0),…, *σ_k_*(*t* = 0)) *when* 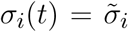 *for any t* > 0}, where 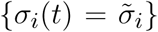 represents the intervention consisting of a specific set of node states. Note that the following two notations are equivalent: 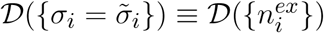. Similarly, we define the logic domain of influence of a node state set, 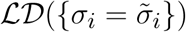, as all the nodes that can be stabilized by the first update in any BFS order-preserving (on the expanded network) update order starting from this given set of node states under all initial conditions. As in the single node state case, the LDOI of a node state set will be a subset of the DOI of the same node state set.

The LDOI of a node state set can be determined by a modified BFS on the expanded network, now using multiple starting points. This does not add complexity to the iterative implementation of BFS: we just need to initialize the queue with the set of given node states. Similar to the case of finding the LDOI of a single node state, we need to deal with the conflicts that may occur during the search process. To be precise, conflict means that during the search we visit a node state that is the negation of a node state included in the intervention. We call such intervention set an incompatible set. The incompatible situation can arise in the following two scenarios. First, a node state in the given set may have a LDOI that was truncated to avoid containing its own negation. The second scenario is when the LDOI of two node states 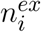 and 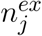 have the property 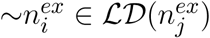 or 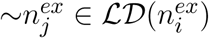, or both. Similar to the truncation we did to find the LDOI of a single node state, we do not include any node state that is the negation of any node state given in the intervention set and we stop searching that branch. We note that this truncation strategy avoids any following conflict. For example, if 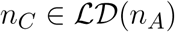 and 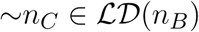, then one may expect that the LDOI of the set {*n_A_*,*n_B_*} will have a conflict between *n_C_* and ~*n_C_*. However, 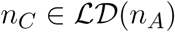 implies that ~*n_C_* requires ~*n_A_*, this means that meeting the conflict between *n_C_* and ~*n_C_*, must be after meeting the conflict between *n_A_* and ~*n_A_*, which is avoided by our truncation strategy.

For a compatible set 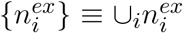, it is guaranteed that 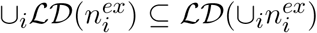. For example, as shown in Fig. 1 (e), the node set {*n*_2_,*n*_4_} is a compatible node set as 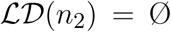, 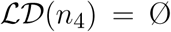 and 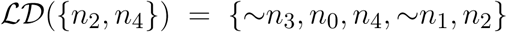. Note 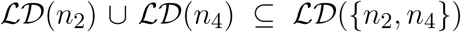. However, for an incompatible set, we just know that the situation 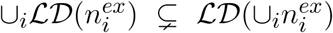 cannot happen and all the remaining situations are possible. In the network of Fig. 1, node set {*n*_2_, ~*n*_4_} is an incompatible node set as 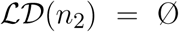, 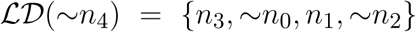 and 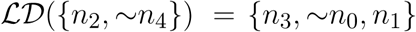. Note that neither *n*_4_ nor ~*n*_2_ are included in 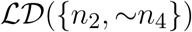 due to the truncation rule and 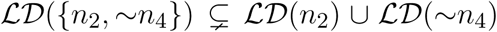. Node set {~*n*_1_,*n*_3_} is another incompatible set as 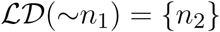, 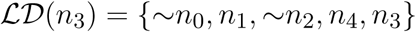 and 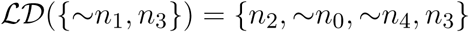. Note that 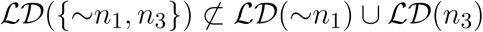 and 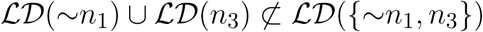.

The properties of the LDOI of a single node can also be generalized to the LDOI of a given node set. For the first key property, let 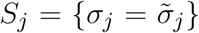 and 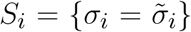 be two sets of node states, if *S_i_* is a subset of any (partial) fixed point and 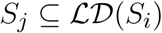, then 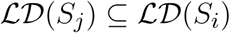. The intuition is similar, the requirement restricting us to consider those nodes which can be stabilized in the long term, that is, we rule out the possibility of *S_i_* being an incompatible node set. For example in Fig. 1 consider *S_i_* = {~*n*_3_} and *S_j_* = {*n*_2_,*n*_4_}. As ~*n*_3_ is part of the stable motif *SM*_2_ = {*n*_0_, ~*n*_1_,*n*_2_, ~*n*_3_,*n*_4_}, corresponding to the fixed point (*σ*_0_,*σ*_1_,*σ*_2_,*σ*_3_,*σ*_4_) = (1, 0, 1, 0, 1), 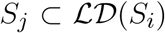 implies 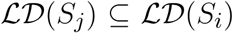. In fact, 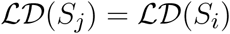.

The second key property also generalizes: if the logic domain of influence of a given node state set contains the set itself, then the logic domain of influence of the set contains at least one stable motif. The intuition and proof is similar to the case of a single node state. Taking the same example, consider *S_i_* = {~*n*_3_} and *S_j_* = {*n*_2_,*n*_4_}, note that both 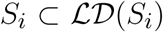 and 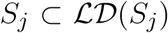, this implies that both 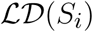 and 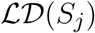 contain a stable motif, which is *SM*_2_ in this case.

Following these examples, we define the core of a stable motif to be a minimal subset of the stable motif whose logic domain of influence contains the stable motif. Here by minimal we mean that no true subset of the core of the stable motif will contain the entire stable motif. The core of a stable motif can be a single node or more than one node. For example, as shown in Fig. 1 (c) ~*n*_3_ is a single-node core of the stable motif *SM*_2_ = {*n*_0_, ~*n*_1_,*n*_2_, ~*n*_3_,*n*_4_}. {*n*_2_,*n*_4_} is another core of the same stable motif as 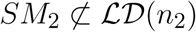, 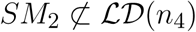 and 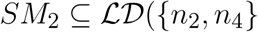.

We also define a driver node (set) of the stable motif to be a node (set) whose domain of influence contains the entire stable motif. The driver node (set) can be inside the stable motif, in which case it is the core of the stable motif; it can also be an upstream node that is sufficient to activate (the core of) the stable motif. We note that stabilization of a stable motif does not require the sustained state of a driver node, that is, oscillations can also lead to the stabilization of a stable motif. An example of this behavior was shown in Fig. 2 (b): node B, which constitutes a self-sustaining stable motif, can stabilize by a single instance of A=1, regardless of the fact that the negative self-regulation of A makes it oscillate.

### 2.7 Target control algorithm

Now that we have equipped ourselves with the tool of LDOI to find the long term effect of a sustained intervention, we can formulate the target control problem as the identification of a node set *S***^*^** whose logic domain of influence contains the target node state set, *i.e.* 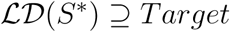. This problem can be framed as a planning search problem [36]. We start with a null set whose LDOI is also null. We repeatedly add a new node to the set until the LDOI of this set contains the target node state set. We use LDOI instead of DOI for this purpose because identification of the DOI is a computationally more difficult problem. Our current solution using LDOI sets a tight upper bound for the optimal solution for the target control problem as 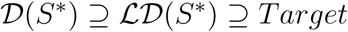.

In order to avoid a full state space search in this combinatorial search problem, we apply a random heuristic algorithm called the greedy randomized adaptive search procedure (GRASP) [37, 38]. The pseudocode is described in Algorithm Table 1 and 2. The algorithm consists of two main phases. The first phase is the construction of a greedy randomized solution and the second phase is a local search to remove any redundancy of the solution.

#### Algorithm 1 GRASP algorithm for Target Control Problem

~~~
1: **procedure** GRASP(*G_expanded*,*Target*,*max_itr*)
2:       *solutions* ← *List*()
3:       **for index** ← 1, *max_itr do*
4:           *solution* ←ConstructGreedyRandomizedSolution(*G_expanded*, *Target*)
5:           *solution* ←LocalSearch(*G_expanded*, *Target*, *solution*)
6:           **if** *solution* **then**
7:              *Solutions.append*(*solution*)
8:           **end if**
9:        **end for**
10:       **return** *solutions*
11: **end procedure**
~~~

#### Algorithm 2 Algorithm for constructing a greedy randomized solution

~~~
1: **procedure** ConstructGreedyRandomizedSolution(*G_expanded*, *Target*)
2:      *solution* ← *Set*()
3:      *α* ← *random*(0, 1)
4:      *candidates* ←Construct_Initial_Candidates(*G_expanded*, *Target*)
5:      **while** *candidates* **do**
6:            *RCL* ←MakeRCL(*candidates*, *alpha*)
7:            *s* ←Select_Candidate(*RCL*)
8:            *solution* ← *solution* ∪{*s*}
9:            **if** *Target* ⊂ *LDOI*(*solution*) **then**
10:             **return solution**
11:      **end if**
12:      Update Candidates(*candidates*)
13:    **end while**
14:    **return Set**()
15: **end procedure**
~~~

In the first phase, we first generate an initial candidate list (line 4 in Algorithm 2). In the simplest case, the initial candidate list is all the non-composite nodes of the expanded network except the nodes in the target set and their negation, both of which are ineligible for control. One can also be more selective to adapt to the specific needs of controlling biological systems. For example, we can forbid the use of certain nodes or node states when constructing the initial candidate list, to incorporate the fact that certain chemical species are harder or even unrealistic to control. Thus these nodes/chemical species will never appear in the final solution since they are not in the initial candidate list.

Then, we begin the procedure of iteratively adding nodes to the trial solution set (which is initially empty) and evaluating whether the LDOI of the trial solution set covers the target set. We form a restricted candidate list (RCL) based on a greedy measure *G*(*v*) defined for each candidate node *v* in the candidate list (line 6 in Algorithm 2). A greedy function is a heuristic score to estimate whether this node should be included in the solution. We discuss several choices of *G*(*v*) below. We determine the minimum score *G_min_* = *min_v_*_∈_*_V_ G*(*v*) and maximum score *G_max_* = *max_v_*_∈_*_V_ G*(*v*) among the heuristic scores of all the nodes. Then we use a previously generated random number *α* from a uniform distribution between 0 and 1 to set a passing score for the RCL as *G_pass_* = *G_min_* + *α* · (*G_max_* − *G_min_*). Then the RCL consist of nodes whose greedy function is no less than the passing score, i.e., *RCL* = {*v* ∈ *V*|*G*(*v*) ≥ *G_pass_*}.

Next we randomly pick a node from the RCL and add it to the current trial solution (line 7 and 8 in Algorithm 2). If after this addition the LDOI of the solution covers the target set, we end the first phase and start the second phase (local search procedure) with this candidate solution (line 9 and 10 in Algorithm 2). Otherwise, we update the candidate node set and start the next iteration toward adding another node from the RCL to the trial solution set. We update the candidate node set by removing the previously added node, its negation and any node in the LDOI of the current trial solution (line 12 in Algorithm 2). We do this latter exclusion because these nodes will stabilize because of the current trial solution, and it is useless to add any stabilized state to the trial solution. We repeat the whole procedure including selecting a node randomly from the candidate set as long as there are still candidate nodes (line 5 in Algorithm 2). We return an empty set if we do not find a solution (line 14 in Algorithm 2).

In the second phase (see the pseudocode in *Supplementary Material* Sec. **1.2**), we start with a candidate solution that covers the target set. We randomize the order of nodes in the candidate solution and then iteratively attempt to remove each node. If after removing this node the LDOI of the modified solution still covers the target set, then we replace the candidate solution with the modified solution. Thus after one iteration of traversing all the nodes, we obtain a final solution. At worst, no node is removed from the set and the final solution is the same as the candidate solution. The randomness in the removal order provides a possibility for obtaining different minimal solutions from the same candidate solution.

In this random heuristic algorithm, we introduce two aspects of randomness in the construction phase, one is the randomness of the passing score by a different *α* for each iteration of solution generation process (line 3 in Algorithm 1) and another is the random selection of a node each time from the RCL inside each solution generation process (line 7 in Algorithm 2). These techniques help strike a balance between the bias of a greedy function and exploring the whole state space [37, 38]. An efficient greedy function/ heuristic score is important to guide the search procedure towards the subspace with the optimal solution. However, a universally efficient greedy function may not exist; rather, the efficiency of a greedy function may depend on the specific network structure and target set. We have implemented five choices of greedy functions *G*(*v*) for a given node state (equivalently, non-composite node of the expanded network): score 1 is the size of the LDOI of that node state (denoted as |*LDOI*|); score 2 is the size of the set of composite nodes which are nearest neighbors of the LDOI of that node state (denoted as |*Comp_LDOI*|); score 3 is a linear combination of the previous two measures with equal weight (denoted as *Scores*_1+2), and score 4 and 5 as the size of the LDOI of that node state with penalty if the LDOI contains a node that is the negation of a node in the target set (denoted as |*LDOI*|_*Pen*1 and |*LDOI*|_*Pen*2). The penalty can be implemented by multiplying this score by −1 (score 4) or by decreasing this score by the size of the largest LDOI among all node states (score 5); both of these implementations ensure that this score becomes non-positive. All relevant code is available at https://github.com/yanggangthu/BooleanDOI.

### 2.8 Computational complexity of the target control algorithm

The time complexity of calculating the LDOI of any set is bounded by *O*(*N_ex_* + *E_ex_*), where *N_ex_* is the number of nodes and *E_ex_* is the number of edges of the expanded network. For each non-composite node in the network, we initially calculate its LDOI and the value of its greedy function, with time complexity *O*(*N*(*N_ex_* + *E_ex_*)), where N is the number of nodes in the original network. We then cache these results to improve the performance of the GRASP algorithm. In the first phase of the GRASP algorithm, we run at most *N* iterations and we need to calculate the LDOI of the trial solution in each iteration, thus the time complexity is bounded by *O*(*N*(*N_ex_* + *E_ex_*)). In the second phase, the time complexity is also bounded by *O*(*N*(*N_ex_* + *E_ex_*)) as we need to go through each node, bounded by *O*(*N*) as a crude estimate, delete the node from the solution and check the modified solution’s LDOI, which is *O*(*N_ex_* + *E_ex_*). The Boolean regulatory functions of biological network models are often nested canalizing rules [39, 40], thus for each node with k regulators there are at most *k* newly generated composite nodes in the expanded network, as well as two corresponding non-composite nodes; each of these nodes have at most k regulators. Thus *N_ex_* is bounded by 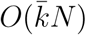, and *E_ex_* is bounded by 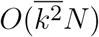. Biological networks are sparse, with an average node in-degree 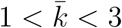 [3]. Thus the complexity of the target control algorithm applied to biological network models is 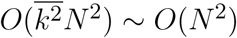 for a well-behaved degree distribution in the sparse limit and bounded by *O*(*N*^3^) for an extremely skewed degree distribution in the sparse limit. Different iterations of the solution generation process (line 3 in Algorithm 1) can be easily parallelized as each iteration is independent.

### 2.9 Damage mitigation as target control

We can generalize the target control algorithm to solve a damage mitigation problem. Consider a Boolean network that has two steady states, one corresponding to the normal state of the system and the other corresponding to a disease state. The system is currently in the normal steady state, but damage to a node, which causes it to stabilize in the opposite state, will lead the system to the disease steady state without any intervention. Under such conditions, previous research has proposed modifying the network topology (as soon as possible, or preventatively) to block the propagation of damage [21]. Here we are interested in designing a damage mitigation strategy to bring the system back to an attractor similar to the normal steady state in the sense that a subset of nodes are in the same state as their states in the normal steady state. This problem is almost the same as the target control problem except that we need to take the permanent damage into consideration. There are two ways of incorporating this. First, we treat this permanent damage as an initial condition and apply network reduction to the system. However, this risks reducing a significant fraction of the nodes in the network, including the target nodes we are interested in. Second, we can apply our GRASP algorithm as above while initializing the solution with the damaged node state(s) and forbidding the damaged node state to be removed in the local search phase in GRASP algorithm. This means that we include the damage as part of the treatment/intervention. When the LDOI of the node state set containing the damage effect covers the target set, the target nodes will stabilize in their desired states after a finite number of time steps under all initial conditions of the subspace of the damaged network. We note that we only need to do this when the damage is a permanent one; when the damage is temporary (i.e. when the node is allowed to go back to its original state), this can be treated as a different initial condition for the target control problem and we can still apply our GRASP algorithm to solve it as DOI/LDOI is robust to any initial condition by definition.

## 3 RESULTS

### 3.1 Application to ensembles of random Boolean networks

We tested the two proposed properties of the LDOI and the target control algorithm on different random Boolean network ensembles. Specifically, we generated an ensemble of 1000 random networks, with size ranging from 15 to 50 nodes and average in-degree ranging from 1 to 2. The Boolean regulatory functions of the random ensemble are required to be effective (irreducible) Boolean functions [41] to be consistent with the generated topology, or nested canalizing functions to simulate biological systems. We have successfully tested and validated the two properties for the LDOI of each node in the generated networks. We also tested and validated the properties of the LDOI of node sets of size up to 3~7 depending on the specific network (as the complexity of testing the property grows faster than *N^k^* for *k* << *N*, where *N* is the network size and *k* is the node set size).

With respect to testing the target control algorithm, we generate 50 random target sets with size 2 or 3 for each random network. It may not always be possible to find a solution for a specific target set for a network, especially when the Boolean network model does not have a (partial) fixed point type of attractor (i.e. if all nodes oscillate in the attractor) or when the desired target state set consists of node states that are part of different attractors, which conflict with each other. In the simulations of the two ensembles mentioned above, we verified that we are able to find a solution for more than 99.5% of the target sets when the target set satisfies two criteria: (i) it is a subset of a (partial) fixed point and (ii) the targets in this set are accessible from nodes outside of this set in the original network (that is, the targets do not consist of source nodes only and do not form a motif without any incoming edges). Note that there can be counter-examples where satisfying these criteria is not sufficient to find a solution. For example, in Fig. 2 (a) and (b), there are no solutions for the target set {~*B*, ~*C*} as the remaining nodes are not enough to activate the composite node in Fig. 2 (b). However, the probability of such situations is small in both random ensembles with moderate size and real biological network. Moreover, the fact that one cannot find a solution through our GRASP algorithm for the target control problem often indicates that the target set is not a reasonable target. It is likely that one would not be able to find a solution in such situation even with a whole state space search.

We also test the performance of different heuristic functions for the target control problem. We calculate the average number of generated solutions for each pair formed by a target set and a network. As shown in Table 1, greedy functions with a penalty for containing the negation of a node state included in the target set (score index 4 and 5) consistently perform better than the greedy functions directly using the size of the LDOI (score index 1 and 3). The intuition behind this is clear, the binary essence of the node state is important and it is thus more efficient to choose from those nodes whose domain of influence does not contain the undesired node state. The second greedy function (|*Comp_LDOI*|) also performs quite well.

**Table 1.** Mean number of solutions found for each target set and random network pair for 50 target sets and 1000 networks. Half of 50 target sets have size two and the other half is of size three; none of them contain source nodes. The 2*^nd^* to 6*^th^* columns correspond to different custom score (greedy function) indexes and notations, which are described in the last paragraph in Sec. 2.7. The second and third row corresponds to the random network ensemble with nested canalizing rules and effective Boolean rules respectively.

| Custom Score Index and Notation | 1<br>$\vert LDOI\vert $ | 2<br>$\vert Comp\_LDOI\vert $ | 3<br>$Scores\_1+2$ | 4<br>$\vert LDOI\vert \_Pen1$ | 5<br>$\vert LDOI\vert \_Pen2$ |
| --- | --- | --- | --- | --- | --- |
| Nested Canalizing Rules | 12.29 | 31.66 | 12.30 | 31.21 | 40.64 |
| Effective Boolean Rules | 26.21 | 61.94 | 26.22 | 57.08 | 66.91 |

### 3.2 Biological Examples

We applied our methodology on four Boolean models of signal transduction networks. In the following we demonstrate our algorithm on two of these, the epithelial-to-mesenchymal transition (EMT) network and the PI3K mutant ER+ breast cancer network. The results on the ABA induced stomatal closure network and the T-LGL leukemia network are shown in *Supplementary Materials* Sec. **3.3** and **3.4**.

#### 3.2.1 EMT network

EMT is a cell fate change involved in embryonic development, which can be reactivated during cancer metastasis [42]. During EMT, epithelial cells lose their original adhesive property, and become mesenchymal cells which leave their primary site, invade neighboring tissue, and migrate to distant sites. A Boolean network model of EMT in the context of hepatocellular carcinoma invasion has been established by Steinway *et al.* [42]. Several predictions of this model were validated experimentally [42, 8]. The EMT network has 70 nodes and 135 edges. The adhesion factor E-cadherin is the sink node; its OFF state indicates the transition to a mesenchymal state. The network has a normal (epithelial) steady state and an abnormal (mesenchymal) steady state. (See details in *Supplementary Materials* Sec. **3.1**). In Fig. 3 we show a simplified version of the EMT network; our analyses were done on the full network.

**Figure 3.**
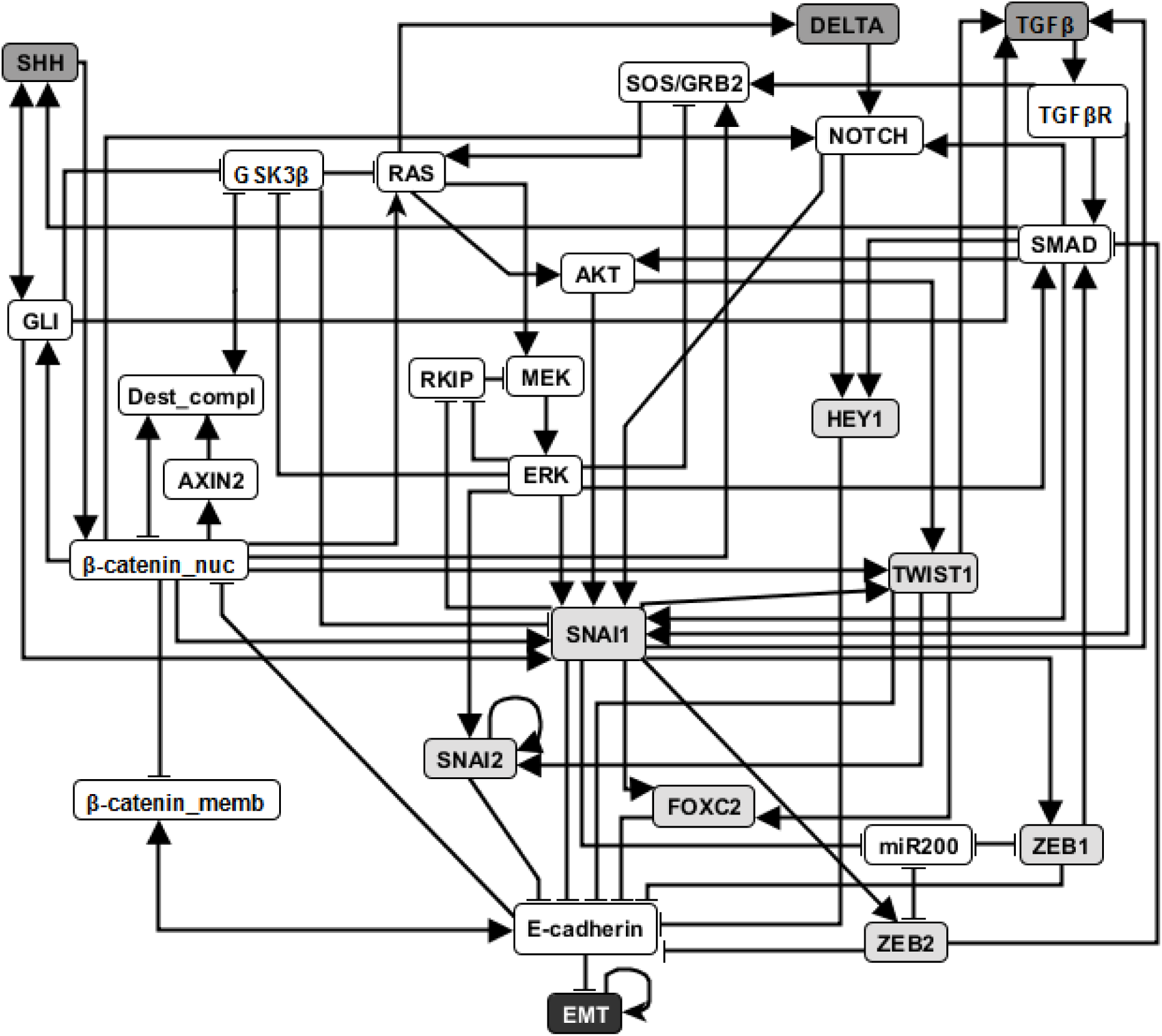
An illustration of the EMT network. Attractor-preserving network reduction was applied to better focus attention on the most relevant nodes. Nodes with light gray background are direct regulators of E-cadherin and nodes with dark gray background represent external signaling molecules. Edges ending with an arrow represent positive regulation while edges end with a flat bar represent negative regulation. See more details in *Supplementary Material* Sec. **3.1**.

Previous research on this network has indicated that sustained activation of TGF*β* signal can trigger EMT through the activation of eight stable motifs [8]. In addition, stabilization of any of these stable motifs can drive EMT. Our target control algorithm shows that any of 60 node states (out of 138 node states for the 69 nodes) can lead to EMT, including the previously established EMT drivers. As we are more interested in designing therapeutic strategies to convert the abnormal steady state into a normal steady state, the negation of EMT is a more relevant target. Previous analysis indicated that when considering an initial epithelial state and turning on the TGF*β* signal, the knockout of any of the transcription factors that downregulate E-cadherin (i.e. knockout of SNAI1, SNAI2, FOXC2, TWIST1, ZEB1, ZEB2, HEY1) or multiple double node knockout combinations (knockout of SMAD and one of RAS, CSL, DELTA, NOTCH, NOTCH_ic, SOS/GRB2) are effective in blocking EMT (i.e. leading to E-cadherin=ON). The effectiveness of transcription factor knockout had been established in the literature; unfortunately these transcription factors cannot be targeted with existing drugs. Several double knockout combinations were validated experimentally in [8] and are more amenable to drug targeting.

For EMT as target, our target control algorithm gives 7 two-node solutions (activation of *β*-catenin_memb and knockout of any of SNAI1, SNAI2, FOXC2, TWIST1, ZEB1, ZEB2, HEY1) and 5 three-node solutions (activation of *β*-catenin_memb, knockout of SMAD and knockout of any of RAS, CSL, DELTA, NOTCH and NOTCH_ic). The main difference between the target control solution and the previously found EMT-blocking single and double knockout interventions is that our target control solution includes the additional control of *β*-catenin_memb. To understand this difference, we note that EMT is in the LDOI of TFG*β*, however, EMT is not in the LDOI of the set consisting of TGF*β* together with any of the previously found EMT-blocking knockout interventions. This indicates that the knockout intervention is effective in the sense that it can block the process of reaching EMT. However, ~EMT is also not in the LDOI of the set of TGF*β* together with any knockout intervention. The knockout intervention is effective when the initial condition is the epithelial steady state, however the knockout intervention does not block EMT for all initial conditions. The target control algorithm, which can block EMT for all initial conditions, requires one more node (*β*-catenin_memb) in the target control solution. In fact, treating this problem as a damage mitigation problem, where the damage is sustained activation of TGF*β*, we verify that EMT is in the LDOI of TGF*β* together with any of the target control solutions.

As established in previous results, the single node EMT-blocking knockouts do not lead back to an epithelial state but rather to hybrid epithelial or mesenchymal steady states [8]. The hybrid epithelial steady state has certain epithelial features, e.g. E-cadherin and *β*-catenin_memb are activated, and also some mesenchymal features, e.g. MEK, ERK and SNAI1 are activated. The hybrid mesenchymal steady state demonstrates the opposite features compared to the epithelial steady state. A good target set to avoid reaching such a hybrid state (which is likely pathological and may even be a worse outcome as the mesenchymal state) would be {~EMT, ~MEK} [8]. The minimum solution found involves controlling three nodes: activation of *β*-catenin_memb, inhibition of SNAI1, inhibition of RAS or RAF. We also find a four-node intervention that does not involve ERK and SNAI1: activation of *β*-catenin_memb, miR2000 and RKIP, and also inhibition of RAS. If the target set is {~EMT, ~MEK, ~SNAI1}, the minimum solution size is found to be six.

Stable motif control indicates that control of at least five nodes is needed to drive any initial state (including the mesenchymal state) to the epithelial state (see Supplemental Table 3 of [8]) Although the control goal is different, one can still see the connection between our target control solution for the target ~EMT and the stable motif control solution (to drive the system to the epithelial state). Specifically, they both require activation of *β*-catenin_memb. Knockout of SNAI1, knockout of TWIST1 or knockout of SMAD and RAS, as one of the target control solutions, also appear as a part of stable motif control solution that does not require control of TGF*β* or TGF*β*R.

These results demonstrate both the accuracy and effectiveness of our target control algorithm, as the solutions found through 1000 iterations are comprehensive (comparable to the solution found through a systematic search of knockout pairs) and indifferent to the distance to the target nodes.

#### 3.2.2 Breast cancer network

Zañudo *et.al.* established a discrete dynamical model of the signal transduction processes involved in the PI3K mutant, estrogen receptor positive (ER+) breast cancer, as shown in Fig. 4 [9]. The model includes 58 nodes, which correspond to proteins, transcripts, drugs, and two cellular outcomes, apoptosis (programmed cell death) and proliferation (cell cycle progression). A fraction of the nodes (16), including the outcome nodes, are characterized by multiple levels, which is implemented by additional virtual nodes, e.g. apoptosis2 corresponds to level 2 of apoptosis, which has a more stringent regulatory function than apoptosis1 (level 1 of apoptosis). This network as implemented is essentially a Boolean network because all the regulatory functions are Boolean [9]. The network model successfully captures the key role of the PI3K/AKT/mTOR signaling pathway in determining the pathological proliferation and survival of cancer cells. In untreated simulated cancers cells, PI3K, MAPK, AKT, mTORC1 and ER signaling are active, leading to high level of proliferation and lack of apoptosis. The network model successfully captures the effectiveness of PI3K inhibiting drugs in leading to low level of proliferation and high level of apoptosis [9]. Through extensive simulations, the network model confirms known drug resistance mechanisms, i.e. additional mutations or other dysregulations that lead to the loss of effectiveness of PI3K-inhibiting drugs. It also predicts new possible resistance mechanisms and the degree of survivability under different resistance mechanisms.[9].

**Figure 4.**
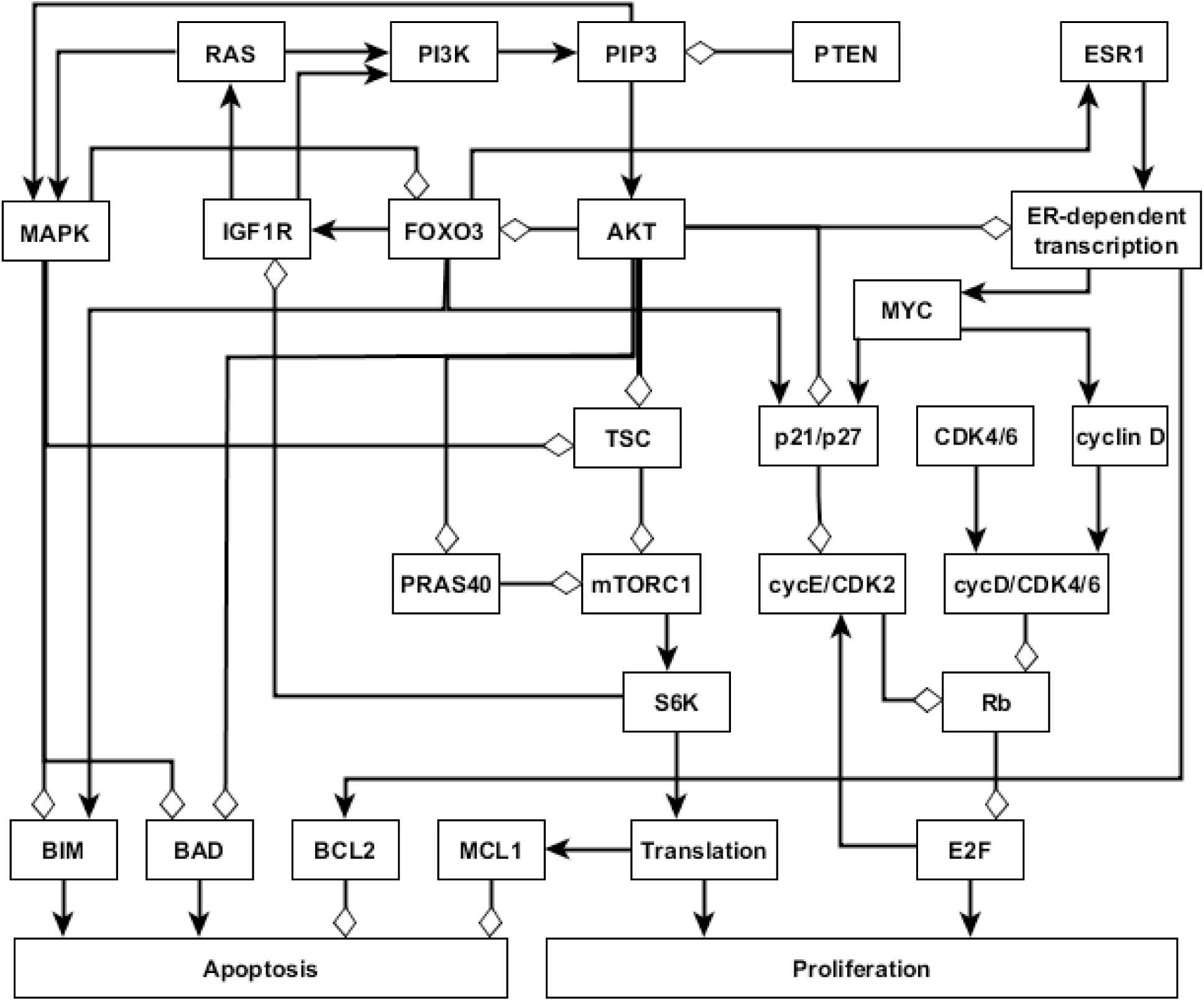
An illustration of the PI3K mutant, ER+ breast cancer network. Attractor-preserving network reduction was applied to focus on the nodes most relevant to our analysis. Nodes are colored according to the signaling pathway that they participate in. Edges ending with an arrow represent positive regulation while edges ending with a hollow diamond represent negative regulation. See more details in *Supplementary Material* Sec. **3.2**

Similar insights can be drawn by applying the target control algorithm to the discrete dynamical network model without doing dynamical simulations, which demonstrate the rich information contained in the network topology and logic and the effectiveness of our control methodology. We obtained a (relatively large) reduced network by considering the system under the relevant initial condition of PI3K mutant, ER+ cancerous state, while keeping the seven drugs as source nodes (see details in *Supplementary Material* Sec. **3.2**.) For example, if we set the target to be high level of apoptosis (Apoptosis = 2), the algorithm’s output is inhibition of PI3K or PIP3. As the target control solution works for any initial condition of the reduced network, this result confirms the key role of PI3K in avoiding apoptosis. If we set the target to be high level of apoptosis and no proliferation, i.e., Target = {Apoptosis2, ~Proliferation}, the algorithm gives multiple two-node interventions as minimal interventions, these consists of either of {~PI3K, ~PIP3} and inhibition of any node in the MYC-CDK4/6 axis of cell-cycle regulation, *i.e.*, {~ESR1, ~ER transcription, ~MYC, ~CDK46, ~cyclinD, ~cycD CDK46, ~Rb, ~E2F}. There are several drugs that can target these nodes. For example, Alpelisib is a PI3K inhibitor, Fulvestrant is a ESR1 inhibitor and Palbociclib is a CDK4/6 inhibitor. This result is consistent with the results found in the [9]: inhibition of PI3K leads to an increase in ER transcriptional regulatory activity, leading to a decrease in proliferation, and simultaneous PI3K and ER inhibition has a synergistic effect in completely blocking proliferation and maintaining a high level of apoptotic activity. If PI3K inhibitor or PIP3 inhibitor is not allowed to be used, the algorithm finds three node solutions involving an AKT inhibitor (e.g. Ipatasertib), MAPK inhibitor (e.g. Trametinib) and inhibition of any node from the MYC-CDK4/6 axis of cell-cycle regulation. In other words, inhibition of AKT together with MAPK provides a similar functionality with inhibition of PI3K. One can also use the LDOI to identify possible drug resistance mechanisms, i.e. perturbations that make PI3K inhibition less effective. As 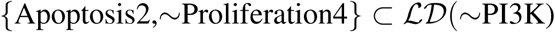, we simply go through all possible two-node interventions containing PI3K inhibitor and screen out those interventions whose LDOI either does not contain Apoptosis2 or contain Proliferation3 or higher level (Proliferation4). We reproduce most of the potential drug resistance mechanism to PI3K inhibitors indicated in Table 3 of [9].

## 4 DISCUSSION

In summary, we have developed the new measures DOI and LDOI to describe the long-term effect of a sustained intervention. We have applied these measures to find solutions to the target control problem in logical network models. This work takes a step forward towards practical control of real biological systems, as illustrated by the applications presented here. The target control solutions we find recover previous predicted interventions obtained by other methods (dynamic simulations and stable motif analysis). As several of these previous predictions are validated experimentally, this agreement also serves as validation of our target control solutions. Notably, by generating a large number of valid target control solutions, we are going significantly beyond previous results. The multitude of predicted target control interventions allows their filtering according to biological or technological considerations.

Here we assumed the existence of a discrete dynamical model. As there are significant uncertainties in the existing models due to the scarcity of experimental information, we estimate the sensitivity of the LDOI measure to the incompleteness of the dynamical model. As the primary way of obtaining causal information that can be used in a logical model is to perform knockout experiments, the predominant causal information indicates a node as being necessary for the activation of another node. For example, if the knockout of either of two regulators A or B leads to a decrease in the activity of target C, we would infer that the logical rule for C is *C* = *A* AND *B*. Suppose that there is a so far undetected regulator of C, which we denote by X. This X will likely also be necessary, which would maintain agreement with the previous observations, *i.e. C* = *A* AND *B* AND *X* is the true rule. Consider the rule for the complementary node ~*C* = ~*A* OR ~*B* in the case of the incomplete system versus the true rule ~*C* =~*A* OR ~*B* OR ~*X*. We can see that the LDOI of any of ~*A*, ~*B*, *A*, *B* will be robust to the addition of X. The LDOI of node *X* and ~*X* need to be established in the true system. The LDOI of node state set {*A*, *B*} will be affected by this change. (However, LDOI of ~*A* and ~*B* will not change.) Thus the size of the solution of the target control problem may increase due to this incomplete information. Due to the binary essence of the Boolean rule, missing a sufficient regulator (an extra OR rule) will give similar results.

The DOI and LDOI can be related to prior research on logical networks. The concept of elementary signaling mode (ESM), originally defined as a minimal subgraph that can propagate a signal from a source node to an output node,[33, 43] can be generalized to start from any node of a directed network and end in any node reachable from it. An ESM on the expanded network is the generalization of a path on a usual directed network. Similarly, the LDOI of a node on the expanded network is analogous to a connected component reachable from a node on a usual directed network. In the same way a connected component reachable from node i consists of nodes that have a path starting from node i, the LDOI of a node consists of all the nodes included in any ESM that starts from that node. Recent work by [44] developed a logic framework to identify causal relationships that are sufficient or necessary. This framework allows an alternative definition of the LDOI. The LDOI of the ON state of a node 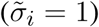 includes all the nodes for which the node is a sufficient activator (these nodes will have 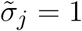) or sufficient inhibitor (these nodes will have 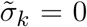). Similarly, the LDOI of the OFF state of a node includes all the nodes for which the node is a necessary activator (these nodes will have 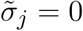) or necessary inhibitor (these nodes will have 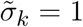).

An algorithm to construct ESMs through a backward search from an output node was presented in [45]; this algorithm can be adapted to find solutions of the target control problem of a single output. If we treat the output node as the root of a backward search, the set of nodes found in the ESM in each search depth (distance from the output node) can serve as a control solution. A truncation technique similar to ours needs to be applied to deal with inconsistent feed-forward or feed-back loops. This algorithm can be generalized to solve the target control problem of a target set by simultaneous search from each target node. We chose to transform the target control problem into a planning search problem; and it has been established that such a planning search problem can be solved in both a forward propagation and a backward propagation approach, or even a mixed approach [36]. It will be an interesting future work if such techniques can improve the efficiency of the algorithm.

This work points out interesting questions as future research directions. First, though evaluating DOI of a node (set) is computationally hard, a better estimation of DOI than LDOI is desirable and can be used to reduce the size of the solution given by our current target control algorithm. Second, the requirement that the solution works for all initial conditions in the setup of the target control problem gives robust solutions, however it may still be conservative for biological systems in certain applications, especially if one is certain about the relevant initial condition subspace. A semi-structural approach (without doing dynamical simulations) to solve the target control problem starting from a subspace of initial conditions are also desirable.

## AUTHOR CONTRIBUTIONS

GY, JZ and RA designed research and methodology; GY and RA performed the analyses; and GY, JZ and RA wrote the paper.

## ACKNOWLEDGMENTS

This research or portions of this research were conducted with Advanced CyberInfrastructure computational resources provided by The Institute for CyberScience at The Pennsylvania State University (http://ics.psu.edu).

## SUPPLEMENTAL DATA

Supplementary Material is attached as required.

